# Latent encoding of movement in primary visual cortex

**DOI:** 10.1101/2024.11.11.623057

**Authors:** Charlie Cosnier-Horeau, Hannah Germaine, Isaiah Choi, Stephen D. Van Hooser, Jérôme Ribot, Jonathan D. Touboul

## Abstract

Neurons in the primary visual cortex (V1) are classically thought to encode spatial features of visual stimuli through simple population codes: each neuron exhibits a preferred orientation and preferred spatial frequency, both of which remaining invariant to other aspects of the visual stimulus. Here, we show that this simple rule does not apply to the representation of major features of stimulus motion, including stimulus direction and temporal frequency (TF). We collected an extensive dataset of cat (of either sex) V1 responses to stimuli covarying in orientation, direction, spatial frequency, and TF to assess the extent of motion selectivity. We show that preferred TF is mostly uniform across the cortical surface. Yet, in over half of V1, the preferred direction is reversed with changing stimulus TF, revealing four distinct map motifs embedded in V1’s functional architecture. Similarly, despite the lack of spatial modulation for the preferred TF map and the lack of invariance for the preferred direction map, we found using convolutional neural networks that direction, TF and stimulus speed can be accurately decoded from V1 responses at all cortical locations. These findings suggest that subtle modulations of V1 activity may convey fine information about stimulus motion, pointing to a novel primary sensory encoding mechanism despite complex co-variation of responses to multiple attributes across V1 neurons.

**Significance Statement:** Understanding how the external world is represented in the brain has long been a central endeavor in neuroscience. In the mammalian primary visual cortex (V1), groups of neurons encode stimulus properties through changes in activity, with “preferred” stimuli eliciting maximal responses, as demonstrated for orientation and spatial frequency. Using an extensive dataset of neuronal responses, we investigated whether V1 also encodes motion speed. We found that cortical response organization remains largely invariant, with neurons exhibiting a uniform preference for temporal frequency. Yet, machine learning algorithms decode motion speed with remarkable precision, revealing that subtle, spatially distributed modulations of activity underlie speed encoding. Moreover, we show that preferred direction flips with speed across 80% of cortex, uncovering a novel co-organization of motion and direction information in V1.

## 1 Introduction

Motion perception, the ability to infer the speed and direction of motion of elements in a visual scene is a central capacity of the mammalian visual system and essential for animal survival. In dynamic visual scenes, motion is characterized by local directions and velocities of points or edges in space, experimentally studied using spatio-temporal gratings characterized by a *speed* (degrees per second) given by the ratio of stimulus temporal frequency (TF, Hz) to spatial frequency (SF, cycles/degree).

The neural mechanisms underlying motion perception have been a central topic of study in neuroscience [Albright and Stoner, 199 The primary visual cortex (V1) has been deemed to have a central role in motion processing since the pioneering works of Hubel and Wiesel [Hubel and Wiesel, 1962] that identified direction-selective neurons in cat V1, later confirmed in primates [Orban et al., 1986, Priebe et al., 2006], cats [Gilbert, 1977, Ohki et al., 2005], ferrets [Moore IV et al., 2005, Weliky et al., 1996, Lempel and Nielsen, 2019, Lempel and Nielsen, 2021], and mice [Andermann et al., 2011].

Local components of motion processed in V1 are then transmitted to higher visual areas, such as the middle temporal area (MT, or V5) in Human or Macaque monkey, for further integration [Dubner and Zeki, 1971, Born and Bradley, 2005, Zeki, 2015, Lempel and Nielsen, 2019]. MT allows for low-speed detection of objects, and evidence in Human suggests that another pathway bypassing V1 brings information directly from the lateral geniculate nucleus for processing fast-moving stimuli [Zihl et al., 1991, Shipp et al., 1994]. Notably, neurons in MT in macaque are highly specialized for motion processing, responding robustly to moving stimuli to produce a coherent representation of complex motion patterns across the visual field. Moreover, these neurons are highly selective for speed [Liu and Newsome, 2003, Lagae et al., 1993, Maunsell and Van Essen, 1983, Perrone and Thiele, 2001], but do not cluster into functional columns with similar speed tuning [Liu and Newsome, 2003].

This speed selectivity of MT neurons contrasts with V1, where experimental data present a more complex picture. Studies reported speed- or TF-tuned neurons in macaque V1 [Orban et al., 1986, Priebe et al., 2006, Foster et al., 1985, Priebe et al., 2006]. Other studies, in human V1, reported that SF and TF preferences cluster spatially in a binary co-representation [Sun et al., 2007]. Similar results have been found in mouse V1 [Ji et al., 2015] or cat [Shoham et al., 1997], where cells also exhibit a continuum from speed to TF selectivity [Andermann et al., 2011], while uniform representation of TF was found in the bush baby (*Otolemur garnetti*) [Khaytin et al., 2007].

To further add complexity to the understanding of SF, TF, and speed tuning in V1, evidence from electrophysiological studies in cats [Saul and Humphrey, 1992] and ferrets [Moore IV et al., 2005] have suggested that while preferred orientation remains invariant to changes in TF, direction selectivity can vary significantly. In ferret V1, it has been shown that direction preference changes with TF, while orientation tuning remains stable across a range of temporal frequencies [Moore IV et al., 2005]. Similarly, in the cat visual cortex, direction selectivity is strongest at low temporal frequencies (1–2 Hz) but diminishes or even reverses at higher temporal frequencies [Saul and Humphrey, 1992]. These findings suggest that TF/speed maps and direction maps could co-vary and should thus be studied together to fully understand how motion is encoded in V1.

In this study, we studied speed selectivity in cat V1 (area 17) using high-resolution intrinsic optical imaging. Leveraging recent advancements in stimulation techniques and analysis methods [Ribot et al., 2013, Sauvage et al., 2017, Kalatsky and Stryker, 2003], we collected an extensive dataset of V1 responses to a variety of stimuli changing in orientation, direction, SF and TF. By combining unbiased computational analyses and targeted selectivity and topology inquiries, we examined how functional maps of orientation and direction varied across 45 combinations of SF and TF (5 SF *×* 9 TF). Additionally, we explored V1’s ability to decode speed, SF and TF using advanced imaging data decoding and statistical learning. We show that spatially distributed V1 activity patterns contain fine local information about the direction and speed of motion. Additionally, while changes in TF left orientation maps unaltered, we found a significant remapping of the direction maps, with smooth variations of the position of the direction fractures, and constant movement speed coincided with fracture immobility. Finally, we investigated topological changes across different speeds and classified these into four distinct topological motifs with varying contributions to speed decoding. This approach reveals how functional maps in V1 integrate motion-related attributes like TF and speed, providing novel insights into the neural encoding of complex motion dynamics.

## Results

### Uniform TF tuning and invariant orientation preference

We began our investigation by examining how changes in stimulus motion properties (direction and TF) affected the reflectance activity in V1. For this purpose, we used a rotating and drifting sinusoidal grating [Kalatsky and Stryker, 2003, Sauvage et al., 2017] to probe responses in cat V1 across all orientations and directions, 9 temporal frequencies (ranging from 0.38 to 5.37 cycles per second) and 5 spatial frequencies (from 0.15 to 1.65 cycles per degree). We computed, for each condition, activity maps reflecting the level of activity of neurons at each given location and for each condition through the recorded hemodynamic signals, averaged over 10 repetitions of the stimulation.

We observed that patterns of activity displayed different responses upon changes in stimulus properties (Figure 1). Because we are interested in stimuli eliciting maximal responses as well as in the spatial reorganization of activity upon changes in stimulus properties, we decompose the signals activity into two independent quantities: (i) the reflectance signal amplitude for a given condition (see Methods), and (ii) the rescaled signal, estimated from the reflectance signal at a given condition by subtracting the lowest value and dividing by the signal amplitude (thus, yielding values between 0 and 1). Although in nature firing rate amplitudes are a function of a variety of properties of the stimulus (e.g., contrast and intensity) and attention mechanisms, it is classically considered that, within the controlled conditions of the experiment, changes in reflectance signal amplitude inform us of the features that elicit maximal firing rates overall. Rescaled signals instead provide information about the spatial reorganization of activity upon changes in stimulus properties. The data shows that varying the stimulus direction (Figure 1A, top row) caused major modifications in the spatial organization of activity, as visible in the rescaled maps but the signal intensity remained constant (right). In contrast, changes in TF did not result in major changes in the spatial organization of activity, but the intensity of the signal showed a clear dependence in TF, with a unimodal profile peaked at 1.53 Hz (Figure 1D).

**Figure 1.**
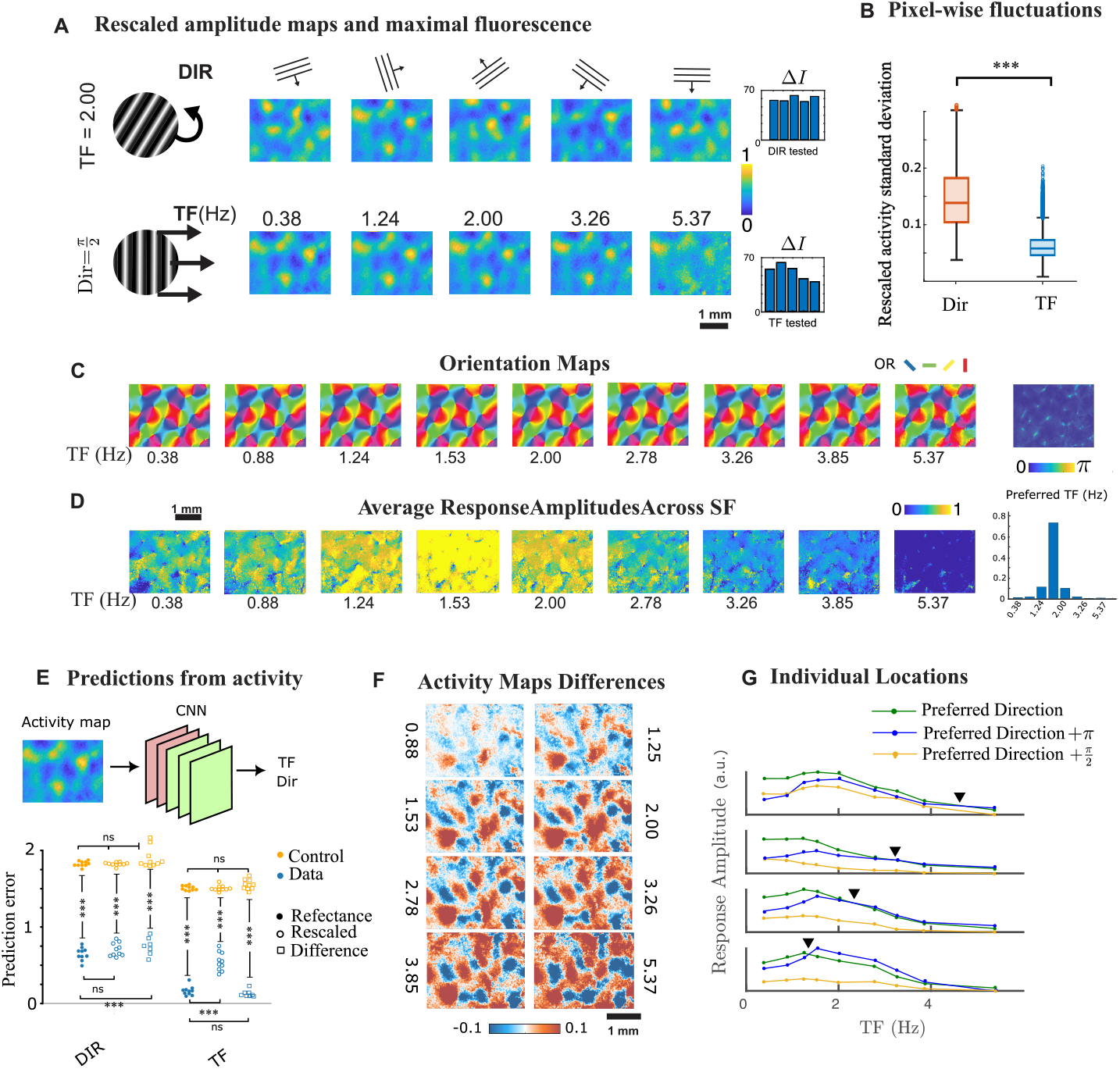
Uniform representation of temporal frequency preference with decodable information. A-D: Invariant features. A. Top: Response amplitude Δ*I* (measured as a change in reflectance, see Methods) for various directions of motion and a fixed TF (2 Hz). Bottom: Response amplitude for various TFs and a fixed direction (rightward, *π/*2) (bottom). In both panels, the stimulus spatial frequency is 0.29 cycles/degree. The responses are rescaled between 0 and 1. The mean response amplitude for each stimulus condition is plotted on the right. B. Standard deviation of the response magnitude at each pixel when varying stimulus direction and stimulus TF. Median standard deviation for direction is 0.11 ± 0.04 mad, while for TF it is 0.03 ± 0.03 mad (two-sample t-test *p <* 0.001). C. Orientation maps are largely invariant to changes in stimulus TF (median standard deviation of 0.2 ± 0.15 mad radian). Right: standard deviation of the response for each pixel, with higher fluctuations near orientation pinwheel centers. Right: Pixel-wise standard deviation of the orientation preference across TF conditions. Higher fluctuations are observed near orientation pinwheel centers. D. Amplitude of responses to changes in TF, averaged across SF at preferred orientation. The preferred TF shows a unimodal distribution (histogram on the right) that peaks at 1.53 Hz, with 90% of locations within the 1.24-2.00 Hz range (cat 2: 81% and cat 3: 78%). E. Top: The architecture of the Convolutional Neural Network (CNN) is composed of five layers (two convolutional in pink and three rectified linear layers in green). This CNN was trained to predict stimulus TF and direction from V1 responses. Bottom: Prediction error (blue dots) shows high predictability for TF and direction compared to controls (same dataset but where TF and Direction were randomly shuffled, yellow dots) (10-fold cross-validation, two-sample t-test *p <* 0.001). F. Difference between V1 response maps between *T F* = 0.38 and other stimulus TFs. These differences show a gradual increase in strength with increasing TFs, as well as smooth variations in spatial organization. Stimulus spatial frequency is *SF* = 0.29 cycles/degree and stimulus direction *π/*2. G. Smoothed responses of four cortical locations separated by 87*µm* along a line, showing changes in response with varying stimulus TF. The preferred direction was determined at TF=0.38 Hz, and response tuning curves are displayed for the preferred direction (green), the opposite direction (blue) and the orthogonal direction (orange). Black triangles indicate switches in preferred direction for each cortical location.

The resulting preferred TF map did not exhibit spatial organization, with a globally spatially uniform dependence in TF and a majority of cortical locations (92%) exhibiting maximal responses for TF in [1.24,2.00] Hz (Figure 1C, and also Figure S1 A-B for two other animals). This uniform preference is of the one found in Bush Baby [Khaytin et al., 2007], and stands in sharp contrast with the spatial organization of other visual attributes, such as orientation or SF, where each cortical location marks a clear preference that spans a wide range of stimulus conditions [Hubel and Wiesel, 1962, Issa et al., 2000, Ribot et al., 2013]. Interestingly, variations in TF had no discernible effect on the representation of orientation (Figure 1D and S1C for two other animals). This map remained strikingly invariant with stimulus TF as quantified by the low median absolute deviation in preferred orientation upon TF variation (averaged across all cortical locations, mad = 0.04 ± 0.02, 0.05 ± 0.03, 0.03 ± 0.02 rad for the three animals (in degrees, 2.29 ± 1.15, 2.86 ± 1.72 and 1.72 ± 1.15), see Figure S1D). Altogether, this data argues for a uniform encoding of TF in V1 and no effect on preferred orientation.

### Smooth Differential Responses encode TF in V1

Despite this apparent uniformity in TF representation, we explored whether changes in activity could convey information about stimulus TF. To infer the presence of information on TF that could possibly be accessible to areas downstream of V1, we trained a convolutional neural network (CNN) to evaluate stimulus direction and TF from reflectance and rescaled responses. Similar approaches were previously applied not only to decode stimuli orientation [Kamitani and Tong, 2005, Haynes and Rees, 2005] and direction [Kamitani and Tong, 2006] in humans, but also to investigate their representation with working memory [Harrison and Tong, 2009, Albers et al., 2013, Xing et al., 2013] and mental imagery [Reddy et al., 2010, Cichy et al., 2012, Stokes et al., 2009, Johnson and Johnson, 2014] and even to classify functional efficiency of aging cortices [Erb et al., 2020]. Moreover, these methods allow for a more detailed assessment of brain-activity patterns, revealing how even non-maximally responding regions contribute to decoding stimuli, as seen in previous studies [Haxby et al., 2001].

Here, we found that the CNN was able to accurately decode both direction and TF from V1 signals with high levels of accuracy(Figure 1E), both from the reflectance signals and from the rescaled activity. The ability to decode the direction of motion from V1 activity is consistent with previous findings [Kamitani and Tong, 2006, Wang et al., 2014, Van Kemenade et al., 2014], and is expected, as strong activity shifts occur in response to changes in stimulus direction. However, given the low level of dependence of V1 response amplitudes on TF, particularly when rescaling maps of activity, it was unexpected to observe that TF could be accurately decoded from brain signals. This suggests that while the TF map is not organized into large-scale spatial patterns preferring various TFs, subtle yet spatially structured variations in the response maps still emerge and may encode TF information in a latent form accessible through advanced decoding methods.

To reveal these possible fluctuations, we computed the difference between rescaled activity maps at a given TF *t*, SF *s* and direction *θ*, noted *A*(*t, s, θ*), and the rescaled activity corresponding to the same SF and direction but a fixed *TF*, here chosen to be *TF*_1_ = 0.38*Hz*. The *difference maps*, defined as *A*(*t, s, θ*) − *A*(*TF*_1_, *s, θ*), lose all the information about the rescaled responses, but revealed a strikingly organized structure that depends on TF, space, SF and direction (see Fig. 1F, as well as Figure S2 for a more exhaustive representation of a set of differential responses as a function of TF and direction or SF). This rich and smooth structure uncovered through this operation, with amplitude and spatial organization showing progressive variation upon changes in TF, reveals a latent encoding of motion in V1. This spatial organization shows that, while the primary structure reflects a strong preference for direction, TF induces second-order fluctuations in the responses that could potentially carry information about the stimulus TF. To test this hypothesis, we trained a CNN to predict TF from the difference maps, and observed that TF was not only decodable from the difference map, but that accuracies was consistent with the reflectance images predictions (Fig. 4E, squares, 2 sided t-test *p* ≈ 0.03), indicating indeed enough information is contained in the differences to decode TF; similarly, we found that we could decode direction from response differences, with a small but significant (2-sided t-test *p <* 0.001) decrease in accuracy compared to when it was decoded from the reflectance data. In Figure S4A, we further compared the learning speed of the CNN when trained on response differences or reflectance images. We observed that learning of TF was significantly faster when CNNs were trained on response differences, with fewer fluctuations and noise in the learning process, suggesting that response differences provided TF information that was more readily accessible compared to the reflectance images.

Going deeper into the structure of the signals, we examined the response amplitudes at four individual cortical locations for multiple directions and temporal frequencies (Figure 1G, smoothed to reveal their structure, see methods). Remarkably, we found that while most directions considered were associated with unimodal responses peaked around the common preferred TF, the dependence in TF of response amplitudes showed distinct profiles between different directions at the same location. In Figure 1G, we depicted responses at the preferred and opposite direction, as well as responses associated with a direction perpendicular to the preferred direction. All three curves showed vastly distinct profiles, which allows the emergence of complex structures in the response difference while maintaining globally uniform preferences in TF and unchanged first-order responses. Furthermore, Figure 1G also confirmed, consistently with previous work [Moore IV et al., 2005, Saul and Humphrey, 1992], that specific locations may change preferred direction according to TF (black triangles in Fig. 1G), an effect we explored in more detail in subsequent sections.

Altogether, these findings show that while we found that the preferred TF map is globally uniform, the minute changes in amplitude with stimulus TF differ from location to location, suggesting that V1 encodes more nuanced information about TF than previously recognized.

### Continuous changes in direction maps fractures

We further explored the structure of changes in direction preference by reconstructing and analyzing preferred direction maps for each combination of temporal and spatial frequency tested. We observed substantial variations in direction maps upon variation of spatial and temporal frequency (Figure 2A and S3 for the other two animals), with a mean absolute deviation tripled compared to that associated with orientation maps (0.12 + 0.11 rad, 0.17 + 0.13 rad, and 0.09 + 0.09 rad for the three animals). These variations in direction maps closely align with changes in direction tuning curves upon TF variation, as suggested by single-cell studies in ferret [Moore IV et al., 2005, Saul and Humphrey, 1992] and exemplified for two cortical locations for different values of stimulus SF and TF (Red and blue curves, Figure 2B). Moreover, the spatial location of these shifts in preferred direction were not confined to the vicinity of pinwheel centers (Figure 2C and S3C for the other two animals). Instead, large regions of the cortical surface switched their preferred direction (Cat 1: 80.23%, cat 2: 59.20%, cat 3: 77.00%), with no significant bias in the preferred orientation of the regions that switch direction preference (Rao test for uniformity, corrected for continuity of the maps; see Methods).

**Figure 2.**
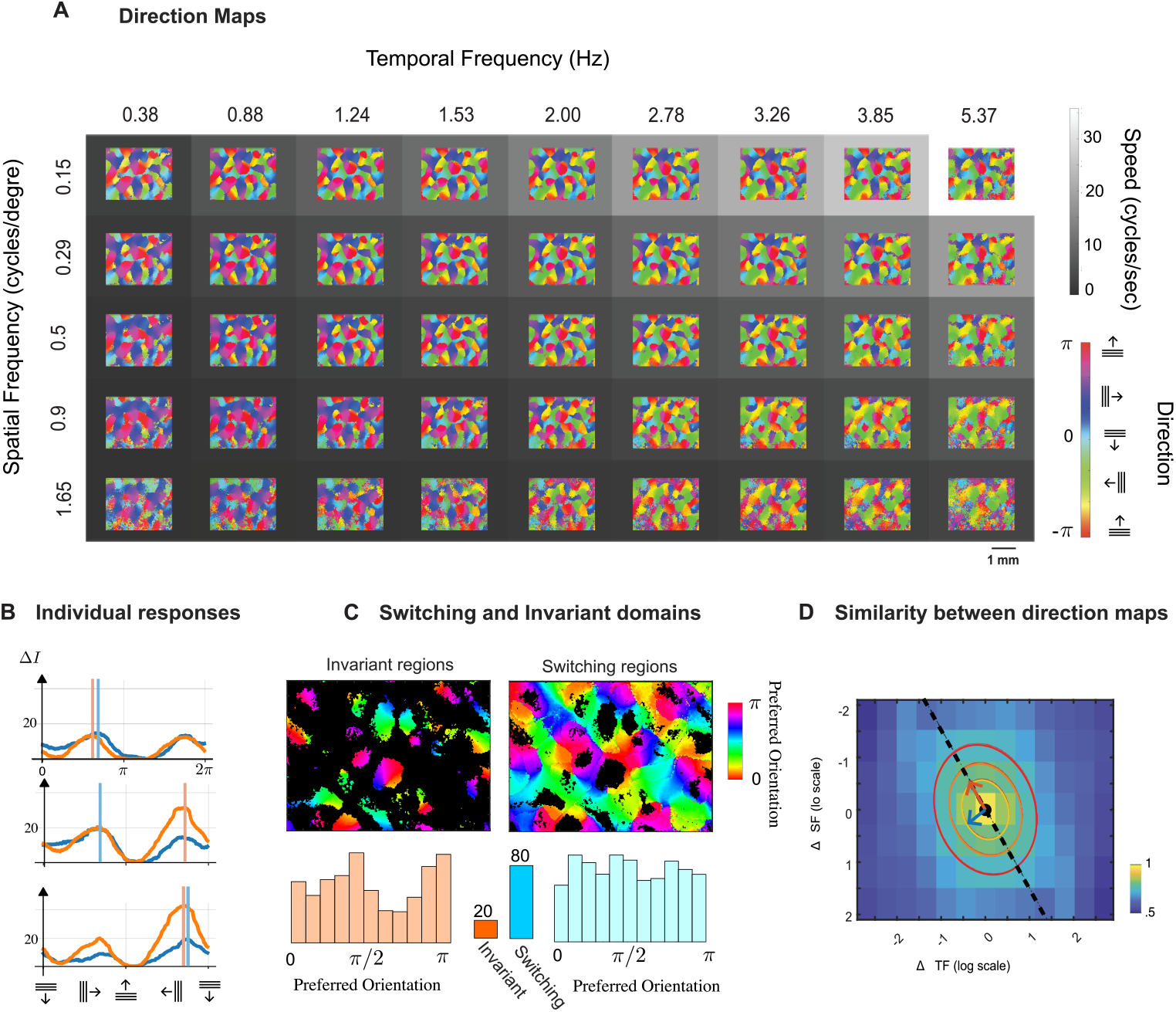
Variations of direction preference representation with stimulus temporal frequency and speed. A. Direction preference maps for all combinations of stimulus spatial frequency and TF,overlaid on a grey scale grid indicating stimulus speed. B Illustration of the direction tuning curves for two different cortical locations (blue and orange curves) and three different conditions (from top to bottom, (SF,TF)=(0.29,0.38), (0.15, 1.24), (0.15,2)). Vertical bars indicate the preferred direction of each tuning curve. C. Representation of cortical locations that do not switch direction preference (left, 19.97% of the cortical area) and those that do switch direction preference (right, 80.03% of the cortical area) overlaid on the invariant preferred orientation map. Histograms of preferred orientation are shown below, and these distributions exhibit no differences when corrected for continuity (*p >* 0.1, see test in Methods). D. Similarity index between pairs of direction maps, plotted according to the logarithmic distance between stimulus TF and spatial frequency. The axis of anisotropy is estimated using a multivariate Gaussian fit, with ellipses representing two level sets of the fitted Gaussian. The main axis is shown as a black dashed line, with a fit of 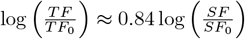. The minor axis is indicated by a blue vector.

The matrix layout of the different maps in Figure 2A additionally suggests that changes in direction maps are not isotropic when stimulus SF or TF are varied independently. Instead, the changes seem dependent on specific combinations of the two stimulus properties. A physically relevant parameter combining SF and TF is the speed. Indeed, when considering the drifting gratings used as stimulus inputs, the speed, expressed as degree of visual angle per second, is defined as the ratio of TF to SF. This speed index is depicted as the greyscale background in Figure 2A. To estimate the axes of maximal sensitivity in direction maps upon changes in SF and TF, we quantified the similarity of direction maps across all conditions. We found that maximal similarity between maps was achieved along a linear axis in the logarithmic scale of TF and SF differences. By fitting a multivariate Gaussian function over a range up to 4.6 octaves of SF and TF differences (i.e. 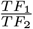 and 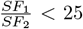), we identified that the main axis of the fit followed the line log(*TF*) ≈ 0.84 log(*SF*) (or *ξ* = 0.84), see Figure 2D. This relationship reflects an intermediate scaling between pure speed and pure TF (cat 2, *ξ* = 0.78; cat 3, *ξ* = 0.94; see Figure S3D). Overall, our findings indicate that there are co-organized changes across the majority of the direction maps as TF and SF vary, following an axis between pure-TF and pure-speed tuning. This suggests that V1 direction maps integrate information from both TF and SF in a coordinated manner, tuned to a combination of speed and TF.

### Fractures remain stable for constant stimulus speed

Modulo *π*, direction preference maps share the typical organization of orientation preference maps, which are continuous except at orientation pinwheels punctual singularities around which all orientations are represented. Due to this constraint, direction maps are bound to display linear singularities, called fractures, where direction preference displays a sharp discontinuity, shifting to the opposite preferred direction (Figure 3A). Topologically, each pinwheel is associated with at least one fracture. Typically, direction fractures consist of linear singularities connecting two pinwheels or forming closed loops.

**Figure 3.**
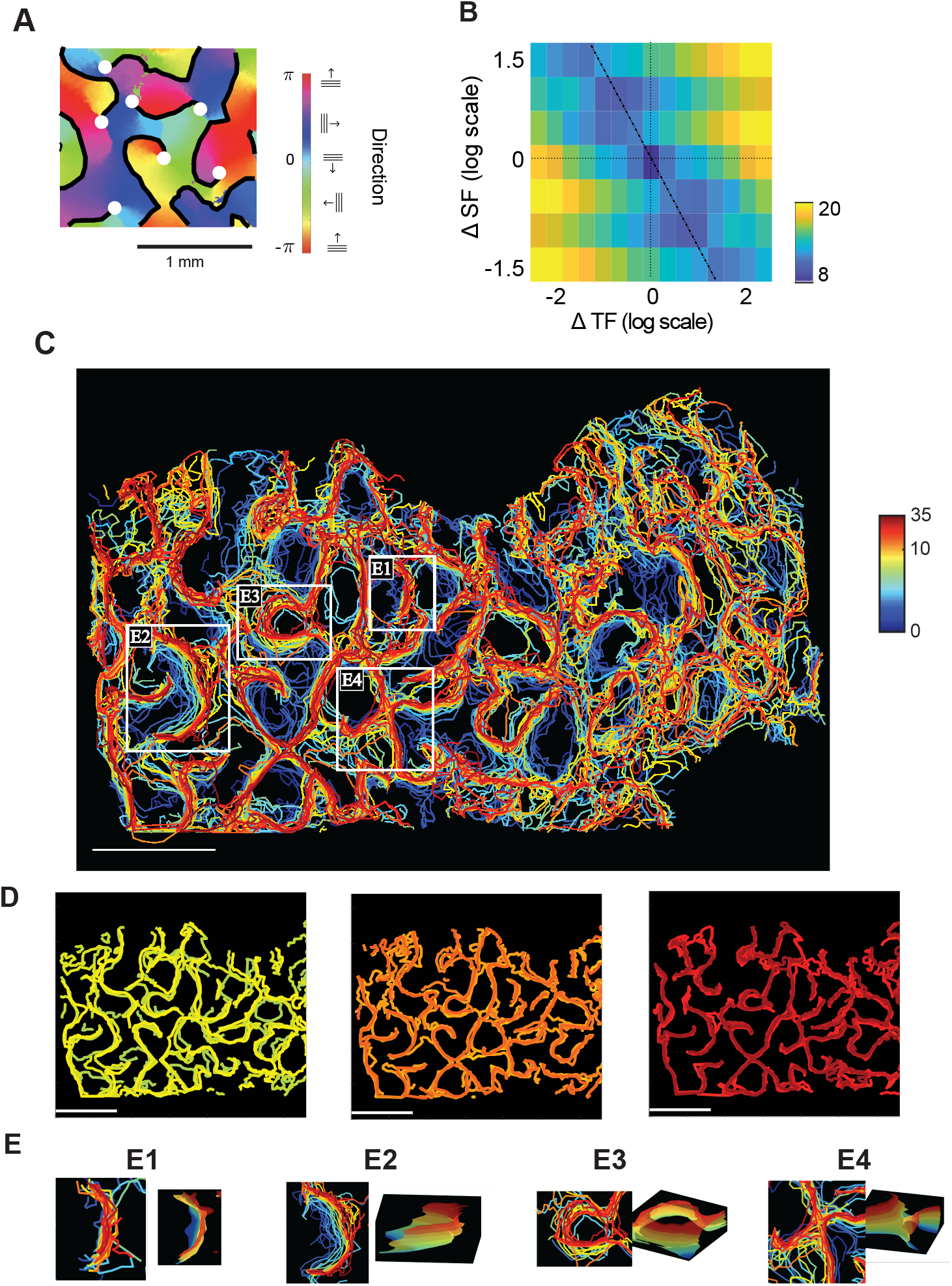
Shifts in direction maps fractures with speed reveal stereotyped motifs. A. Illustration of preferred direction map fractures (black contours) overlaid with orientation pinwheel centers (white dots) for *TF* = 1.53 Hz and SF=0.29 cpd. B. Average distance between fractures in pairs of direction maps, plotted according to the logarithmic difference between stimulus TF and spatial frequency. Minimal changes occur along a slanted axis (dotted line). C. Direction fractures computed for each combination of stimulus TF and spatial frequency, colored-coded according to speed (in logarithmic scale). D. Representation of direction fractures across different speed intervals, where minimal changes occur as shown in B: left panel shows speeds in [2.45, 5.5] degree/second (7 conditions), middle panels in *v* ∈ [6, 14] degree/second (7 conditions), and right panel in *v* ∈ [14, 33] degree/second (5 conditions). E. Illustration of typical fracture movements with speed, shown as a zoomed-in view from C with a 3D surface representation where the (*x, y*)-axes represent the cortical surface and (*z*-axis) represents speed. E1: Invariant direction fractures. E2: Traveling fractures, where fractures shift in one direction with speed. E3: Bubbling fractures, where one fracture splits into two, with one forming a closed loop. E4: Crossings fractures, where two fractures exchange their connections.

To investigate changes in direction preference and potential topological changes in direction maps, we meticulously tracked direction fractures in each of the 45 direction maps (9 TFs *×* 5 SFs, Fig. 3C and Figure S3A for the other two animals). These fractures exhibited continuous variations in response to changes in speed, covering large regions of the cortical map. To quantify changes in fracture positions, we computed the cumulative distance between fractures (see Methods) as a function of stimulus conditions. Similarly to the observations in Figure 2D, when plotted against the logarithmic SF and TF differences (Figure 2B), we observed that the greatest invariance occurred along a slanted axis as expected from our results in Figure 2D. This invariance became even more apparent when fractures were grouped according to speed: positions of fractures showed only minor variations for stimuli associated with similar speeds, even when SF and TF differed significantly (Figure 3D).

As the stimulus speed changed, some fractures remained spatially stable (Figure 3E1), while some others were gradually shifting, in a majority of cases moving along a given axis maintaining the identity of the pinwheels they are connected to (Figure 3E2). Occasionally, fractures formed detached closed loops (bubbling, Figure 3E3), or exchanged which pinwheel they were connected to (saddles, Figure 3E4). Of note, similar instances were found in all animals considered (see Figure S3B, right). These topological changes actually arise from the continuous morphing of the fracture shapes, as illustrated in the three dimensional reconstructions of the fracture surfaces, where speed is represented as altitude. Altogether, these findings reveal that direction fractures in V1 display major topological modifications upon variation of stimulus speed, resulting in reversals of preferred direction across large portions of the cortex, but remain largely stable for stimuli with similar speeds.

### Local decoding of speed from V1 activity

The observation that direction maps are not unique, but vary with speed, raises questions about the possibility of decoding the direction of stimuli with distinct speeds, as well as the independent decoding of speed and direction. To explore this, we used a similar convolutional neural network as in Fig. 1E) and trained a regression algorithm to predict speed from reflectance or rescaled data (Figure 4A). Remarkably, we found that reflectance (Figure 4B) or rescaled (Figure 4C) brain activity images produced highly accurate speed and direction predictions, with performance significantly better than control datasets where the association between brain activity images and speed was randomized. To assess whether this decoding performance required a deep or complex network, we tested multiple CNN architectures with varying levels of complexity (Figure S4B). All tested architectures, including those with minimal convolutional and linear layers, were able to extract stimulus information significantly above chance, confirming the robustness of the observation of decodability of speed from the activity maps, irrespective of the specific parameters of the algorithm used. Furthermore, we found that speed and direction information can be decoded locally, in cortical regions as small as 123 *×* 123*µm*^2^ (10-by-10 pixels with our resolution) (Figure 4C). Even in these small regions, the decoding accuracies were comparable to that of larger areas, suggesting that speed encoding is a local property of V1 responses. This may imply the animal’s capacity to detect complex movements associated with various speeds and directions at different locations in the visual scene.

**Figure 4.**
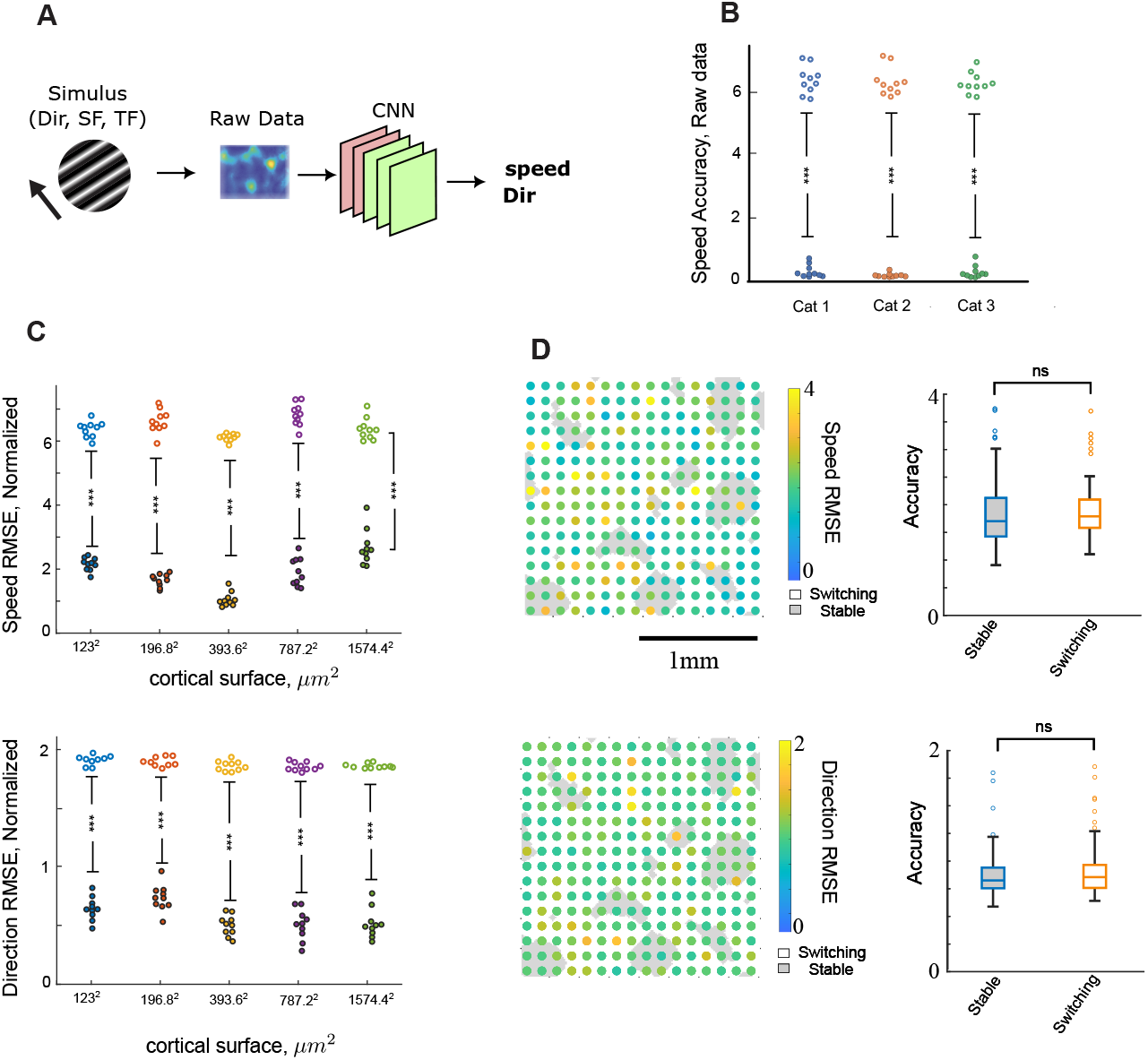
Preferred direction and preferred speed are accurately decoded from local V1 responses despite uniform or variable representation. A. Illustration of the convolutional neural network architecture. It is composed of five layers: two convolutional layers (pink) and three linear layers (green), with rectified linear activation functions. B. Prediction error for speed accuracy across three animals shows high predictability compared to controls (same dataset but where TF and Direction were randomly shuffled) (10-fold cross-validation, two-sample t-test *p <* 0.001). C. Prediction error for speed accuracy (top) and direction accuracy (bottom) across cortical regions with different size (from 123×123*µm*^2^ to 1574.4 *×* 1574.4*µm*^2^), shows high predicability compared to controls (same dataset but where TF and Direction were randomly shuffled) (10-fold cross-validation, two-sample t-test *p <* 0.001). D. Spatial organization of coding accuracy for preferred speed (top) and preferred direction (bottom) of non-overlapping cortical windows with 10 *×* 10 pixels (123 *×* 123*µm*^2^, colored circles) overlaid with invariant (grey) and switching (white) direction domains. Right panels show no significant difference in coding accuracy between invariant and switching direction domains (2-sample t-test *p >* 0.1).

Finding that speed was decodable at local scales thus allowed the investigation of potential correlations between cortical position in V1 and speed coding accuracy. To do that, we computed on a fine grid of non-overlapping regions of 123*x*123*µm*^2^, speed and direction decoding accuracy over a large area of V1 (Figure 4D, left panels, top for speed and bottom for direction). While there were some spatial fluctuations in accuracy, no clear correlation emerged between these spatial fluctuations and changes in preferred direction with stimulus speed. To quantify this observation, we labeled each window as either “switching” or “stable” depending on whether the majority of pixels contained in that area changed direction preference with speed. Notably, we found no significant difference in decoding accuracy between these two regions (Figure 4D, right panels). This suggests that the spatial variations in direction preference and speed decoding accuracy are two sides of the same coin, the differential changes in response amplitudes rather than consequences of one another. Also, no correlations were found between the coding accuracies in speed and direction.

## Discussion

Our findings provide new insights into how motion attributes are represented in V1 and their potential implications for visual perception. We demonstrated that TF preference is uniformly distributed across V1, lacking spatial modulation typical of other visual attributes. Moreover, direction preference exhibits significant remapping in response to changes in TF and speed, with about 60-80% of the cortical area switching preferred direction under varying conditions, putting in perspective the notion of a well-defined direction map in V1, but revealing a collection of maps smoothly varying upon changes in TF. In sharp contrast, the OR map was invariant to changes in TF, revealing that some maps are robust to changes in a given attribute while others are variable. Despite this variation in direction maps and uniform TF representation, the changes in cortical activity associated can provide fine and local information about the stimulus, as we showed using convolutional neural networks (CNNs) to decode motion attributes from brain signals, including TF, speed, and direction. These results formally establish that maps organized according to stimulus preferences covering the range of percepts, or even representations of visual attributes that are invariant upon changes in other stimulus properties, are not necessary for visual perception. This offers a novel perspective on how V1 may encode motion attributes.

Pioneering investigations of TF representation in cat V1 have reported the presence of binary domains with a corepresentation of SF: regions responding preferentially to low SF and high TF or high SF and low TF [Shoham et al., 1997]. While subsequent studies have revealed that SF representation in cat V1 spans a broad range of preferences and is predominantly continuous [Issa et al., 2000, Ribot et al., 2013], the case of TF representation has not been reexamined. Here, we demonstrate that TF preference in V1 is mostly uniform across the cortical surface, with a 80-90% of cortical locations exhibiting maximal responses in the 1.24-2 Hz range, and lacking the spatial modulation suggested by earlier studies. These findings align closely with results obtained in promisian primates Bush Babies [Khaytin et al., 2007], where TF preference was similarly reported to be uniformly distributed, centered around 2 Hz. Additionally, TF had a significant impact on the representation of direction preference. An organization for direction preference in cat V1 has been reported [Weliky et al., 1996, Swindale et al., 2003, Ohki et al., 2005], but these studies examined responses to a single TF of stimulation. Our data reveal that direction map is not invariant with respect to stimulus TF and that TF and direction should be studied together. Interestingly, these shifts in direction preference with stimulus TF align with specific map motifs, suggesting an underlying structure to this variability.

Our results also shed new light on the uniform coverage hypothesis, a foundational principle of cortical organization [Swindale et al., 2000] postulating that the different attributes are encoded into parallel maps that favor even representations of combinations of attributes [Hübener et al., 1997, Yu et al., 2005, Nauhaus et al., 2012]. Two of our findings challenge the uniform coverage hypothesis for motion-related features: (i) the predominantly uniform distribution of TF preference contradicts the expectation of even representation across the attribute’s full range, and (ii) the dynamic changes in direction preference with stimulus TF violate the principle of invariant representation. These observations suggest that uniform coverage may not extend to motion-related features and that alternative mechanisms might govern their cortical representation.

Our CNN analysis provides compelling evidence that functional maps in cat V1 are not necessary for the perception of motion attributes. Indeed, despite the lack of a structured TF map and the variations of direction maps, we demonstrated that TF, speed, and direction information could be accurately decoded from local V1 responses at any cortical location. Notably, no differences in decoding accuracy were found between zones with invariant direction preference and those with switching direction preference. The question of the functional significance of maps for visual perception echoes findings in squirrel monkeys, where, variability in the expression of ocular dominance columns -from well-developed to nearly absent- does not impair binocular vision [Horton and Adams, 2005, Adams and Horton, 2006], challenging the presumed necessity of these maps for stereopsis. Similarly, in rodents [Girman et al., 1999, Ohki et al., 2005], lagomorphs [Murphy and Berman, 1979] and grey squirrels [Van Hooser et al., 2005], robust visual processing occurs without the columnar organization of orientation or direction, underscoring that maps are not indispensable for perception. Instead, we suggest that distributed activity patterns in V1 may convey rich information about motion attributes.

How, developmentally, these co-representations arise remains a central question. Mathematical or computational models, capable of successfully reproducing different aspects of V1 response properties [Miller, 2003, Miller, 2016, Ferster and Miller, 2000 Kaschube et al., 2008, Chariker et al., 2021, Antolík et al., 2024, Ben-Yishai et al., 1995, Teich and Qian, 2006], may collectively suggest that plasticity and the nature of visual stimuli underlie map organizations. In particular, models based on the statistics of natural images and their processing [Olshausen and Field, 1996, LeCun, 2012, Gregor and LeCun, 2010, Antolik and Bednar, 2011, Zhu and Rozell, 2013] or neurogeometric models [Petitot, 2017, Montobbio et al., 2020, Citti and Sarti, 2 tend to suggest that geometric structures and the nature of stimuli are the source of functional architectures. An intriguing avenue of research would be to explore these models for speed invariance, fractures. Some of our topological observations and rules of change arise naturally from biological constraints, such as the co-organization of orientation and direction tuning, and stability of orientation tuning across TF changes. Testing if models of V1 and visual perception generate similar responses to varying TF and speeds offers an interesting perspective of this work.

Consistent with physiological perception, the stimulation we used was binocular, which does not allow inferring whether the structures observed emerge from binocular interactions or if they are the result of the superposition of the input received by both eyes. Further experiments will be necessary to test this by stimulating one eye at a time, which would require adapting the experimental protocol to accommodate possible longer experimental times.

Intrinsic optical imaging signals are indirect measures of neuronal activity, reflecting the impact of neural spiking on localized changes in blood oxygenation and volume. Because it provides access to mesoscopic signals over relatively large cortical areas for extended periods, it is well-suited for large-scale mapping of functional organization. Its spatial and temporal resolution is, however, limited, and typically prevents inferring precise patterns of single-cell activity. A significant advance was made in understanding the relationship between hemodynamic signals and neural activity, particularly in the physiology of neurovascular coupling that underlies intrinsic optical imaging and functional magnetic resonance imaging (fMRI). Fine experiments highlighted the correlation between spiking vascular activity [O’Herron et al., 2016] and even revealed the existence of orientation selectivity at the level of individual blood vessels in cat V1, but also that spiking and hemodynamic signals are partially decoupled. Moreover, the consistency between hemodynamic signals and spiking activity depends on the cortical layer and on independent anatomical constraints, such as vessel anatomical orientation through the cortex and capillary organization. Signals were also found to be transported through these vessels to distant regions of the cortex [Cho et al., 2022, Martineau et al., 2024]. All of these observations are fundamental limits to the resolution at which hemodynamic signals can reflect local spiking activity. This motivates combining, in future studies, optical imaging techniques with single-cell resolution techniques to further understand how spiking activity supports the changes in responses we reported, and to assess consistency between neuronal responses and speed selectivity at the single-cell level.

The combination of beautiful and rule-bound topological changes with the co-organization of response properties to all four stimulus properties (OR, DIR, TF, SF), naturally leads to developmental and structural questions. Developmentally, we know that direction selectivity requires visual experience in carnivores[Li et al., 2006, Li et al., 2008] and primates[Hatta et al., 1998], so it is of interest whether co-organization with speed tuning, in particular, follows a strict developmental ordering or is influenced by experience. Further, direction maps and orientation maps are co-organized spatially in columns about pinwheel centers, and given the finding that direction maps change with TF /speed and orientation maps remain constant a natural question arises of whether there is a hierarchy of priority and stability in V1 feature processing, with a strong robustness of orientation tuning invariance, and whether this stability coincides with bottom-up versus top-down activity modulation. The prior segregation of motion processing to higher-order regions, and stationary processing to V1 followed a bottom-up theory of building complexity in processing, but the recurrent connections from these “higher-order regions” in visual processing back to V1 calls into question the extent of the observed motion information arising as a consequence of “top-down” information transfer [Lempel and Nielsen, 2021]. One approach to addressing this question is investigating direction map changes under conditions of targeted optogenetic/chemogenetic silencing of projections to V1 from regions downstream.

## Material and Methods

### Surgical procedure

All experiments were performed in accordance with the relevant French guidelines and regulations. Intrinsic optical imaging was applied on three young adult cats of either sex in good health and with no apparent malformations or pathologies. Animals were anesthetized with saffan (initial dose, 1.2 mg/kg, i.m.; supplements, 1:1 in saline, administered intravenously as needed). Following tracheal and venous cannulation, electrocardiogram, temperature, and expired CO2 probes were placed for continuous monitoring and animals were secured in the Horsley-Clarke stereotactic frame in preparation for acute recordings. The scalp was incised in the sagittal plane, and a large craniotomy was performed over area 17. The nictitating membranes were then retracted with eye drops (Neo-Synephrine 5%, Ciba Vision Ophthalmics), and the pupils were dilated with atropine eye drops (atropine 1%, MSD-Chibret). Scleral lenses were placed to protect the cornea and focus the eyes on the tangent screen 28.5 cm away. Animals were then paralyzed with an infusion of Pavulon (0.2 mL/kg (0.4 mg/kg), i.v.), and breathing was assisted artificially through a tracheal cannula with a 3:2 mixture of N2O and O2 containing 0.5-1.0% isoflurane. The respiration frequency was adjusted to 18 breaths/minute, and the volume adapted to the ratio of exhaled CO2 (partial pressure of carbon dioxide at 4%). Paralysis was maintained throughout the recording by continuous infusion of a mixture of Pavulon (0.1 mL/kg/h) diluted in glucose (5%) and NaCl (0.9g/L). Following the recording session, animals were sacrificed using a lethal dose of pentobarbital.

### Optical imaging

The cortex was illuminated at 545 nm to reveal the vascular pattern of the cortical surface, and at 700 nm to record the intrinsic signals. The focal plane was adjusted to 500 µm below the cortical surface. The optic discs were plotted by tapetal reflection on a sheet of paper covering the tangent screen placed 28.5 cm in front of the animal. The center of the screen was situated 8 cm below the middle of the two optic discs. Intrinsic optical signals were recorded while the animals were exposed to visual stimuli displayed on a CRT monitor subtending a visual angle of around 75^°^ *×* 56^°^. Frames were acquired by CCD video camera (1M60, DALSA) at the rate of 40 frames per second and were stored after binning by 2×2 pixels spatially and by 12 frames temporally using the LongDaq system (Optical Imaging). Images were acquired with a resolution of 12.3 *µ*m/pixel.

### Stimulation

Full-screen visual stimuli were presented continuously to the animal. Each stimulus consisted of sine-wave gratings drifting in one direction and rotating in a counter-clockwise manner [Kalatsky and Stryker, 2003, Sauvage et al., 2017]. The angular speed of rotation was one cycle per minute. The contrast was set at 50% to ensure the production of smooth sine wave gratings [Xu et al., 2007]. Nine temporal frequencies ranging in a logarithmic scale from 0.38 to 5.87 Hz, and 5 spatial frequencies ranging in a logarithmic scale from 0.15 to 1.65 cycles/degree were presented in random order. Ten full rotations were presented for each pair of temporal and spatial frequencies. Stimuli were presented binocularly to avoid extending the total acquisition time (∼ 7 hours) and to maintain the physiological stability of the preparation. The center of rotation of the grating was fixed at the center of the visual display. The center of rotation of the stimulus was the center of the visual display. The impact of the stimulus rotation was minimal relative to the range of grating drift speeds tested: its impact, increasing with eccentricity, reached only speeds of ∼ 4.9^°^*/s* at the corners of the display and thus may produce a slight shift in speed correspondence away from the central region.

### Image processing

To ensure that the data is recorded from area 17, we relied on a functional criterion based on spatial frequency sensitivity: data from A18 were rejected from further analysis based on responses evoked by low spatial frequency stimuli (0.15 cpd, optimal in A18) versus higher spatial frequency stimuli (0.5 cpd, optimal in A17).

Data analysis was based on the method developed by Kalatsky and Stryker [Kalatsky and Stryker, 2003, Sauvage et al., 2017] to extract periodic signals from intrinsic data using Fourier decompositions. For each SF and TF, data were preprocessed to remove slow-varying components using a multivariate analysis technique, the generalized indicator function method [Yokoo et al., 2001, Ribot et al., 2006]. A key limitation of intrinsic optical imaging is that the hemodynamic response acts as a low-pass filter on functional signals [Sauvage et al., 2017]. To correct for the delays and amplitude attenuation associated with this filtering, we applied a frequency-domain inverse filter based on a first-order low-pass model with a 5-second time constant [Sauvage et al., 2017]. This correction step is crucial for accurately recovering stimulus-locked neural activity. A low-pass spatial filter (with a Gaussian kernel of 5×5 pixels and a 2-pixel standard deviation) was also applied to smooth the data. These operations resulted, for each SF and TF, in maps of the changes in reflectance according to direction, noted Δ*I*. To smooth individual pixel responses in Figure 1G, we used a median kernel on SF, TF and direction, where the median was taken on neighboring TF and SF, and neighboring direction in a cone of 25 degrees of opening.

To calculate orientation and direction maps, we first extracted the Fourier coefficients at the frequency of stimulation (fundamental frequency, or first harmonic H1) and the second harmonic (H2) of the angular component of the response at each pixel, for each SF/TF pair. Preferred orientation was defined as the phase of H2 [Kalatsky and Stryker, 2003]. By contrast, direction preference requires more careful analysis, as the phase of H1 alone is insufficient and known to yield biased estimates [Sauvage et al., 2017, Swindale et al., 2003]. To overcome this, we reconstructed the direction tuning curve at each pixel by summing the tuning curves derived from both H1 and H2 components. Preferred direction at each pixel was then defined as the stimulus direction that maximized the resulting combined response. This approach yielded direction maps that closely matched those obtained using the standard technique based on episodic stimulation followed by Von Mises fitting [Sauvage et al., 2017].

The similarity index between two direction maps used in Figure 2D was estimated as follows. Consider two direction maps *θ*_1_(*x, y*) and *θ*_2_(*x, y*), where (*x, y*) are the coordinates on the cortex and that are associated to the same orientation map (that is, *θ*_1_(*x, y*) − *θ*_2_(*x, y*) = 0mod*π*, is defined by the average number of pixels whose preferred direction differ by more than *π/*2:

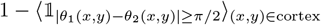

where ⟨·⟩_(*x,y*)∈cortex_ denotes the average on all points of the cortical map, and 𝟙_*condition*_ denotes the indicator function, equal to 1 when condition is satisfied and 0 otherwise. The choices of an indicator function and of a threshold *π/*2 are used to emphasize changes in the map that do not depend sensitively on the threshold. A test using the natural distance, ⟨|*θ*_1_(*x*) − *θ*_2_(*x*)|⟩_*x*∈cortex_, provides similar qualitative results.

### Fracture Detection

To identify these fractures, we used an automatic extraction, identifying the maximal gradient in direction within a single map across x and y coordinates. As shown in Figure 3A, fractures tend to start and end at orientation pinwheel centers, which topographically makes sense given the co-organization of orientation and direction preference maps. Given the changes seen in direction maps in Figure 2, we proceeded to quantify the difference in fractures as a function of changing TF and SF in order to compare the axis of change.

### Convolutional Neural Network

All Convolutional Neural Networks were implemented in Python 3 using PyTorch 1.12.1. Unless specified otherwise, they were trained on cropped images of size 256×256 pixels (i.e., 3.23 *×* 3.23*µm*^2^, for computational efficiency. All neural networks used shared the same architecture, composed of the following operations:

- `Conv2d(1, 6, 5) + RELU`: Convolutional step with 6 filters, that takes as input the reflectance, rescaled or difference *N × N* pixels data (these are single-channel images, thus the input size is 1) with *N* = 256 unless specified otherwise. The size of the convolutional kernels is 5 pixels (63*µm*). 6 filters are computed, resulting in 6 (*N* − 4) *×* (*N* − 4) images. The results were rectified using RELU functions.
- `MaxPool2d(2,2)` reduces the dimension by taking, for each pixel, the maximum in a window of size 2 *×* 2 (yielding 6 output images of size (*N/*2 − 3) *×* (*N/*2 − 3))
- `Conv2d(6, 16, 3) + RELU`: the resulting 6 images are then subject to another 2D convolution with kernel size 3 *×* 3, and 16 filters, rectified with a RELU function. This results in 16 (*N/*2 − 5) *×* (*N/*2 − 5) images.
- `AdaptiveAvgPool2d(1,1)`takes each of the 16 outcomes and returns their average. This yields a 16-dimensional vector.
- `Linear(16,120) + RELU + Linear(120,84) + RELU + Linear (84,1)` the 16-dimensional vector is then processed through a series of 3 linear layers serving as a feedforward network to yield the regression.

Loss was considered to be mean squared error, made periodic when predicting direction. The optimizer was Adams, learning rate 0.001 by default (0.0005 for windows smaller than 20 *×* 20 pixels, or 255.2 *×* 255.2*µm*^2^). All results reported here are associated with such a regression algorithm. Classification for TF (where the output is a tensor of size 9 instead of 1 and loss is a traditional categorical classification accuracy associated with the percentage of accurate classification) yielded similar results. For smaller windows of size less than 20 *×* 20 pixels, we replaced the size of convolutional kernels by smaller kernels of size 3 *×* 3 pixels. Batch sizes were set to 32, and the number of epochs was set to 200, although convergence happened after around 100 epoch or less depending on the window size.

We used 10-fold cross-validation to generate our results. Controls were generated by using identical processes but shuffling the labels to be learned. Comparisons between the dataset and the controls were done using two-sample t-test.

Figure S4B reports accuracies for simpler CNN architectures, described below:

**Model 0** `Conv2d(1, 20, 5) + RELU + MaxPool(2,2)`

`+ AdaptiveAvgPool2d(1,1) + Linear(20,1)`:

Model 0 is an elementary network based upon the same operations as the reference model above but with a single convolutional layer and a single linear layer. For more power, it also increased the number of convolutional layers to 20 instead of 6.

**Model 1** `Conv2d(1, 20, 5) + RELU + MaxPool(2,2)`

`+ AdaptiveAvgPool2d(1,1) + Linear(20,30)+ RELU+ Linear(30,1)`

Model 1 is similar to Model 0, except that it has 2 linear layers for the regression, with dimension of the first layer’s input (and second layer’s output) 30.

**Model 2** `Conv2d(1, 10, 5) + RELU + MaxPool(2,2)`

`+ AdaptiveAvgPool2d(1,1) + Linear(10,30)+RELU+`

`Linear(30,50)+RELU+ Linear(50,1)`: This model now builds upon Model 2 and add a third linear layer, with output dimension of the second layer increased to 50.

**Model 3** `Conv2d(1, 6, 5) + RELU + MaxPool(2,2)`

`+ Conv2d(6, 10, 5) + RELU + AdaptiveAvgPool2d(1,1)`

`+ Linear(10,1)` Model 3 returns to two convolutional layers as the reference model (but with the second convolution having 10 filters) and reduces the number of linear layers to 1.)

**Model 4** Conv2d(1, 6, 5) + RELU + MaxPool(2,2) Conv2d(6,16,3)+RELU + AdaptiveAvgPool2d(1,1) +

Linear(16, 120)+RELU+ Linear(120,1) This model includes the same two convolutional layers as the reference model, but reduces the number of linear layers to 2, with the output dimension of the first layer equal to 120.

## Author Contributions

JDT and JR designed the research, the analytic tools, analyzed the data and wrote the paper. JR collected and processed the data. CCH and HG performed the analyses and wrote the paper. HG analyzed fracture displacement. IC ran machine learning codes to measure information processing. SVH designed the study and wrote the paper.

## Supplementary Material

**Figure S1:**
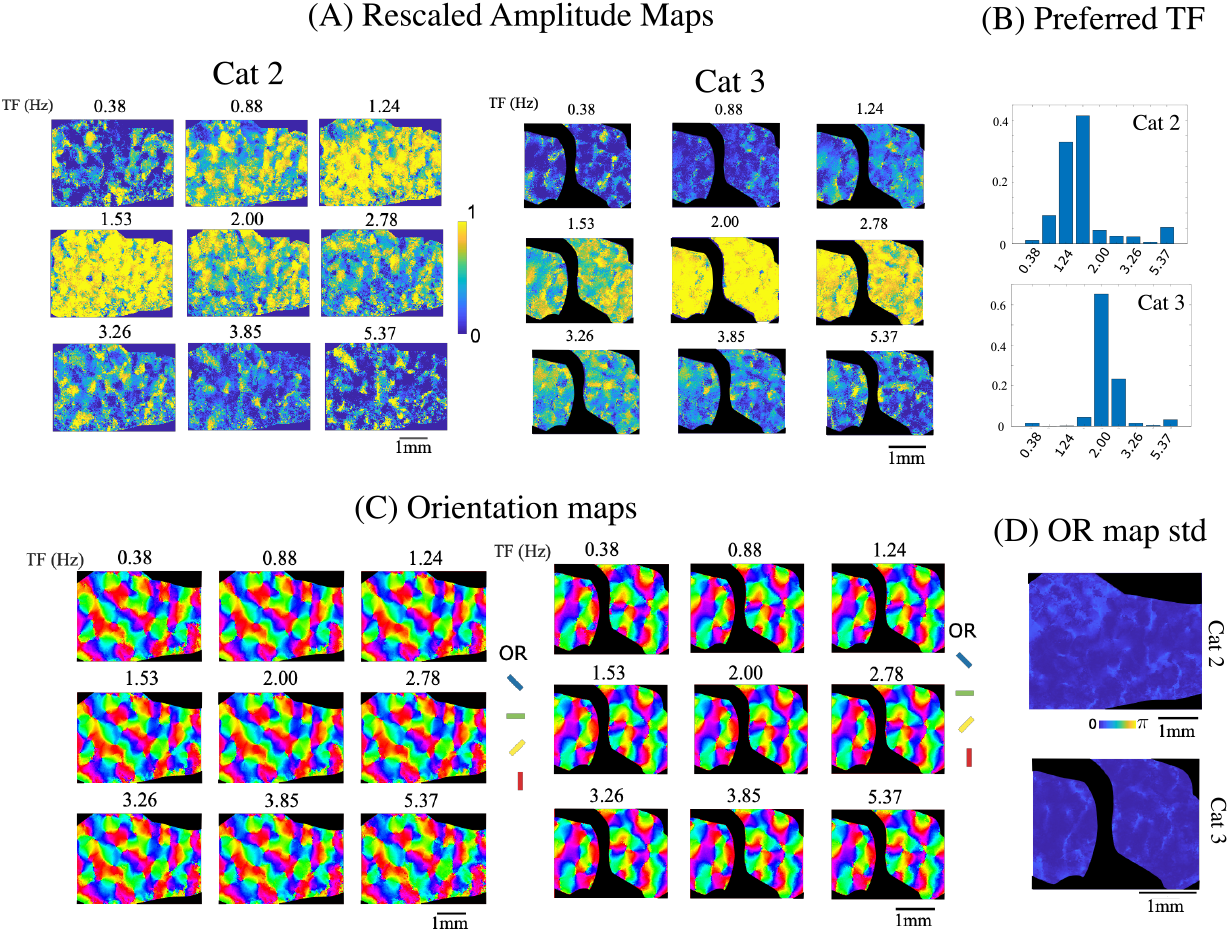
Optical imaging responses in cats 2 and 3 (A) Rescaled signals as a function of TF and (B) histogram of preferred TF (analogous to Fig. 1C), (C) Orientation maps as a function of TF and (D) their pixel-wise standard deviation (C-D compare to Fig.1D). This confirms the observations made in cat 1 of a uniform representation of TF in V1 with a peaked distribution and an overall invariance of orientation maps as a function of TF.

**Figure S2:**
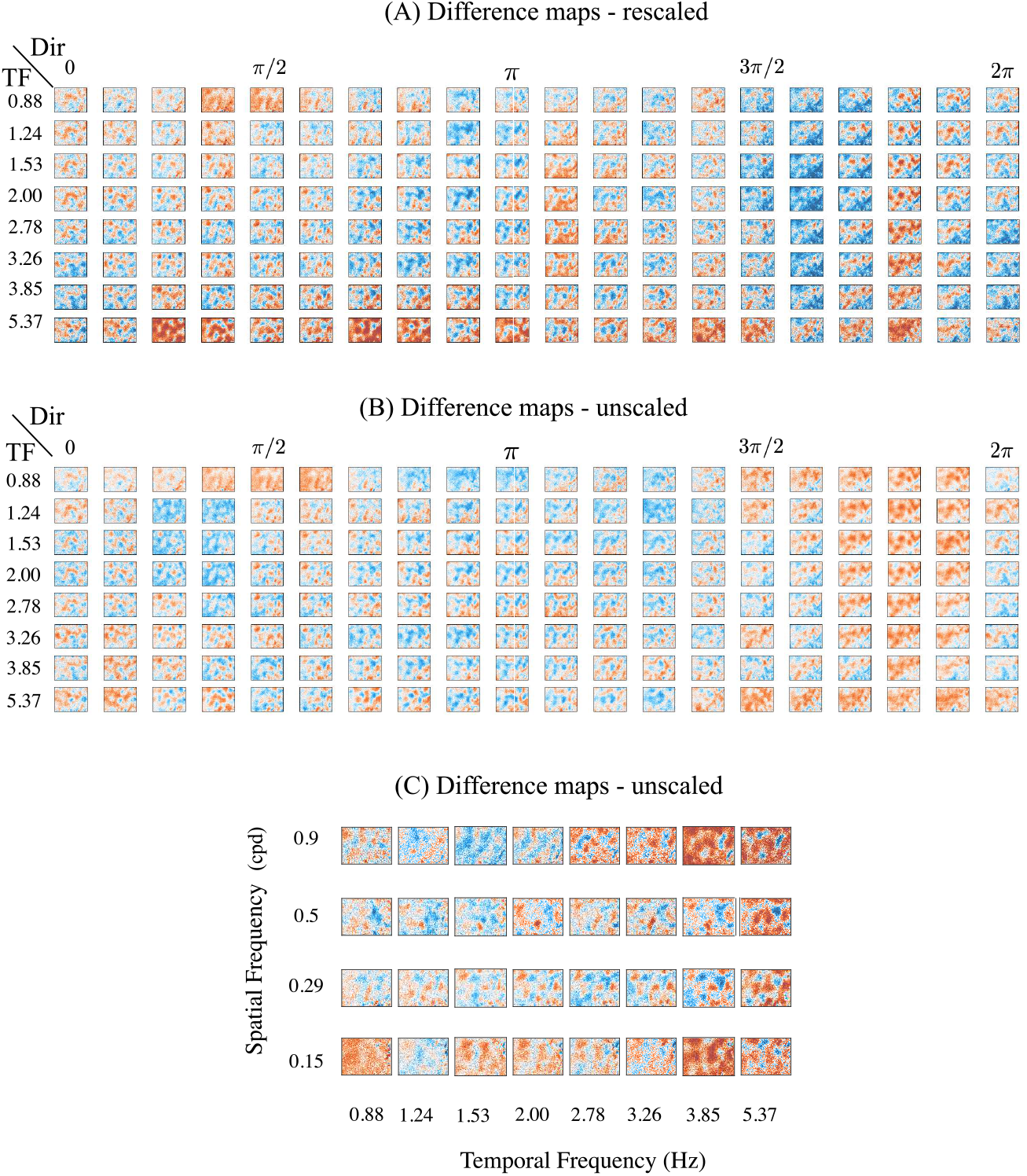
(A-B) Difference activity maps for various directions and temporal frequencies (*SF* = 0.29 cycles/degree), with a reference map of *TF* = 0.38 (scalebar from dark blue to dark red with, for A: a fixed difference scale clipped within [− 0.5, 0.5] and white for 0), and for B: same data, but each figure is individually scaled in [0,1]. The colorbar grows from 0 to 1, with 0.5 corresponding to white. C. Difference maps as a function of SF and TF with fixed direction *π/*2 rad, represented with a common scalebar clipped at [− 0.2, 0.2]. Similar to Figure 1F, we observe that the difference maps are highly structured and reflect fine changes in activity upon variation of TF, SF and direction.

**Figure S3:**
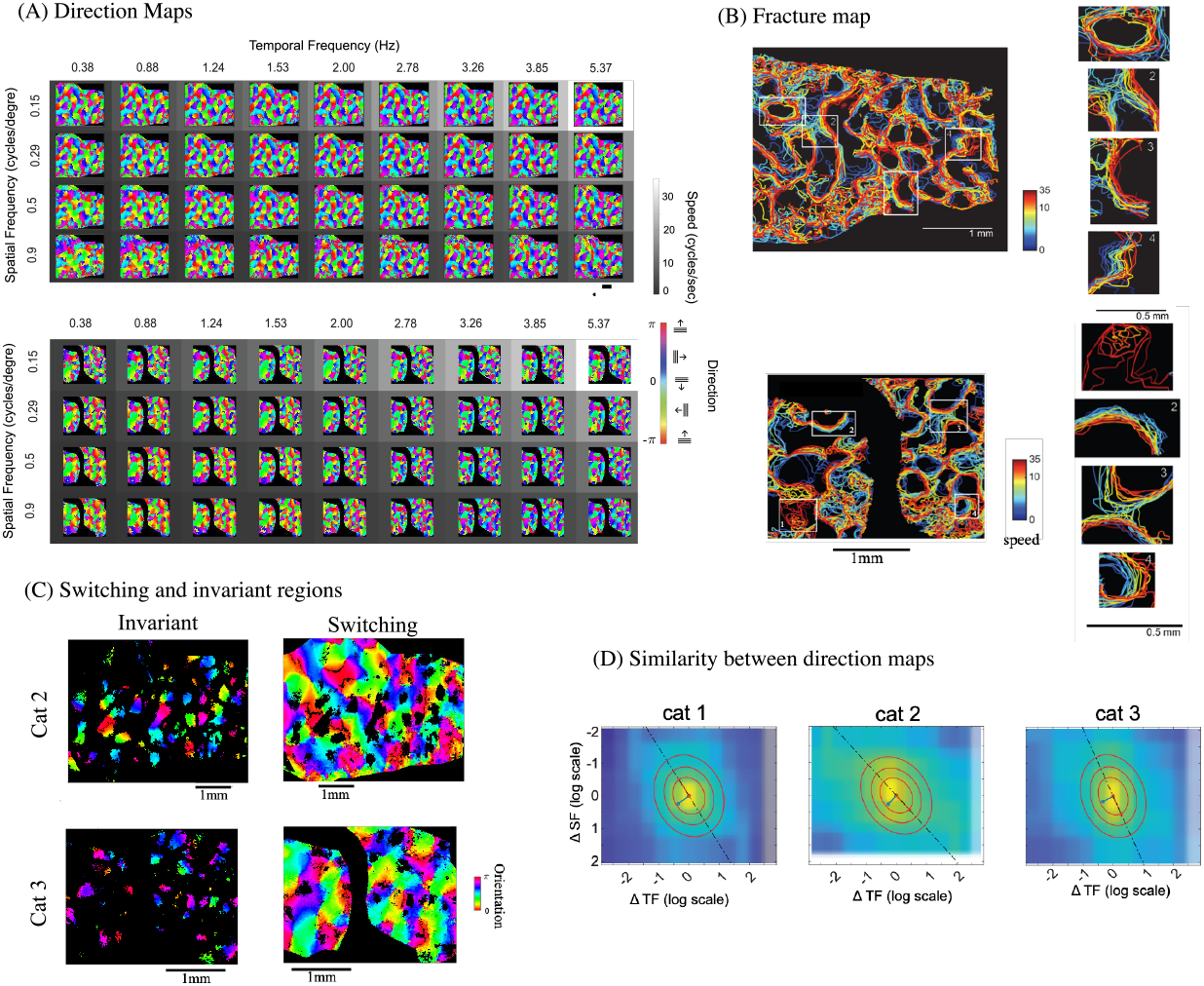
Changes in direction preference, cats 2 and 3. (A) Changes in direction preference as a function of TF and SF for cats 2 and 1 (for cat 1, see Figure 2A) and (B) location of the fractures as a function of speed (color) as well as 4 examples of the types of fracture movement (for cat 1, see Figure 3C and E). (C) represents the preferred directions in the invariant or switching regions (Figure 2 C for cat 1 analyses). (D) represents the similarity between direction maps as a function of the logarithmic difference in SF and TF, with a Gaussian fit (ellipses and vectors) in logarithmic axes within 2.5 octaves. Significant linear relationships with respective slopes 0.84, 0.78 and 0.94 were found, with a ratio of eccentricities *λ*_1_*/λ*_2_ of 1.63, 1.47 and 1.52.

**Figure S4:**
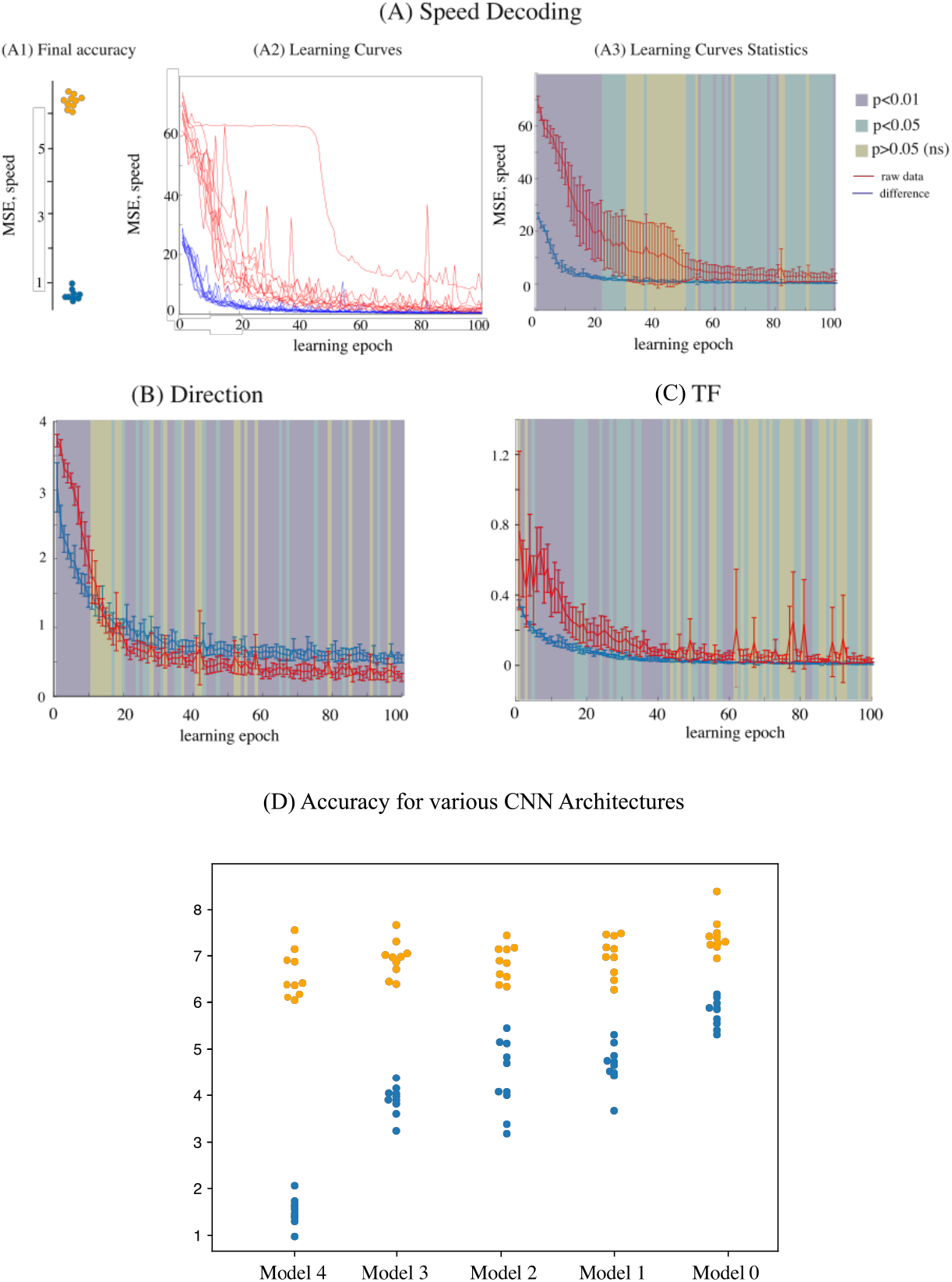
Stimulus decoding. (A) Accuracy of speed decoding for a CNN trained with difference maps. (A1) Significant learning is achieved compared to control (10 fold cross validation, 2-sided t-test *p <* 0.001). (A2) Show validation accuracy as a function of learning epochs (32 learning steps) for the CNN trained with difference maps (blue) or with the reflectance maps (red) for each of the 10 folds of training. Learning from difference maps is faster and more consistent, while learning from the reflectance maps is slower, more variable, and subject to local minima. (A3) shows a statistical analysis of these curves (mean ± sem), with the p-value level associated to a 2-sample t-test for each epoch plotted in color in the background. We can see a significant difference at initial epochs, confirming the fact that learning is significantly faster when training is done for the difference maps, but final accuracies show less difference, and, at the final epoch, no significant difference is found. (B) and (C) are analogous to (A3), but correspond respectively to the learning of direction and TF. Similar, yet less pronounced, phenomena happen for learning of direction and TF, with however a significant degradation of learning direction when training with difference data. (D) Speed decoding accuracies on 5 models with increasing degrees of complexity (see Methods for description of the Models). Blue: decoding accuracy (MSE), compared to control (orange). All models show significant learning (*p <* 0.001). Data used: cat1.

